# From individuals to ecosystems: carrion beetle responses to heatwaves impact nutrient cycling

**DOI:** 10.64898/2026.10.13.758632

**Authors:** Isobel Grieve, Arthur Broadbent, Sheena Cotter, Ainoa Pravia, Natalie Pilakouta

## Abstract

Climate extremes, such as heatwaves, are becoming more frequent and severe, leading to significant disruptions in key ecosystem processes. The direct effects of heatwaves on these processes have thus received increasing attention, yet the indirect effects of heatwaves, as mediated through animal behavioural responses, are still unknown. Insects play a fundamental role in carrion decomposition by modifying carcasses and regulating nutrient transfer to soils. Understanding how insect responses to climate extremes scale up to ecosystem processes is therefore crucial. In this study, we used a widespread carrion beetle, the burying beetle *Nicrophorus vespilloides*, to experimentally investigate the direct and indirect effects of a simulated heatwave on carrion decomposition dynamics, and the consequences for soil nutrient cycling. We found that burying beetle presence increased carcass mass loss by 355% under ambient temperatures, and that this was accompanied by significant increases in dissolved organic carbon (+86%), ammonium (+71%), and nitrate (+47%) concentrations. However, the positive effects of burying beetle presence on carcass mass loss were severely diminished by heat stress. Path analyses revealed that this was related to marked reductions in certain beetle behaviours, such as carcass preparation, as well as a reduction in brood size. These behavioural and reproductive shifts were, in turn, linked to decreased soil nitrate concentrations and soil moisture. Together, these findings demonstrate that the indirect effects of heatwaves on decomposition processes, as mediated by shifts in the behaviour and traits of animal ecosystem engineers, can be as important as their direct effects. Such indirect effects must therefore be incorporated into our understanding of how key ecosystem processes respond to climate extremes.

## Introduction

Climate change is causing rapid and severe alterations of ecosystem structure and function (Nunez *et al*., 2016; Rull, 2021). Increasing global mean temperature and heatwave events may breach species thermal thresholds and alter life history and behavioural strategies (Hughes, 2000; Weiskopf *et al*., 2020). This is particularly true for ectotherms such as invertebrates due to their limited capacity for thermoregulation (González-Tokman *et al*., 2020; Stocker *et al*., 2024). As invertebrates are fundamentally important for the functioning of terrestrial ecosystems, these environmental stressors are likely to affect ecosystem processes both directly and indirectly via changes to invertebrate thermal and behavioural ecology (Prather *et al*., 2013; Cardoso *et al*., 2020). However, our understanding of the relative importance of direct and indirect effects of climate change, as mediated by changes in invertebrate behaviour, on key ecosystem processes remains limited (Wu *et al*., 2021).

Organic matter decomposition is a fundamental ecosystem process that is central to nutrient and energy cycling (Swift *et al*., 1676; Barton *et al*., 2016) and closely linked to the activities of scavengers (Wall *et al*., 2008; Luan *et al*., 2021; Tuo *et al*., 2024). However, most research on decomposition has focused on plant detritus (Benbow *et al*., 2016), and not animal carrion, which is also an important organic matter source. Of the research investigating carrion decomposition, most has been in the context of forensic entomology and not ecosystem functioning (Carter *et al*., 2007; Newsome *et al*., 2021; Patterson *et al.,* 2022). Yet, carrion can be disproportionately important for nutrient cycling (Barton *et al*., 2013; Keenan *et al*., 2018) as it forms ‘islands of fertility’ that decompose up to three times faster than plant detritus and contributes to community heterogeneity and landscape complexity (Swift *et al*., 1676; Carter, *et al*., 2007). As such, both the direct and indirect effects of global climate change on carrion decomposition could have serious consequences for wider ecosystem functioning (Stephenson *et al*., 2025).

Invertebrates are among the principal drivers of carrion decomposition (Payne, 1665; Barton and Evans, 2017). Much of this influence may arise through behaviours that extend beyond the direct consumption of carrion tissue, yet most studies focus only on the presence of decomposers, rather than behavioural drivers (Wu *et al*., 2021; McIntyre *et al*., 2024). Evidence from dung beetles, woodlice, and ants show that behavioural strategies can have substantial effects on decomposition and nutrient cycling, particularly when they fully monopolise a detrital resource (deCastro-Arrazola *et al*., 2020; Torabian *et al*., 2024). For example, dung manipulation and digging capability were positively related to dung removal and subsequent seedling emergence (deCastro-Arrazola *et al*., 2020). Furthermore, increased sheltering time associated with desiccation stress decreased litter breakdown by woodlice (Leclercq-Dransart *et al*., 2016), and the effect of ants (*Formica subsericea)* on leaf litter decomposition was reduced under heat stress, likely mediated by shorter ant surface activity (Del Toro *et al*., 2015).

Burying beetles (Coleoptera: Staphylinidae: *Nicrophorus*) are important drivers of carrion decomposition in many Palaearctic ecosystems (Hoermann *et al*., 2018), as they locate and bury small vertebrate carcasses to complete their life cycle (Scott, 1668; Potticary et al., 2024). By burying carcasses, *Nicrophorus* beetles effectively monopolise the carcass, excluding many other scavenger guilds (Trumbo, 2023). Once buried, a breeding pair of burying beetles and their brood can consume a small vertebrate carcass almost entirely within two weeks (Scott, 1668; Ilardi *et al*., 2021). Such behavioural control over carcass access has important implications for decomposition dynamics. For instance, breeding burying beetles can markedly alter carrion-derived nutrient cycling: temporarily increasing soil organic matter (Ilardi *et al.,* 2021), and significantly increasing nitrate, phosphorous and potassium concentrations (Hoback *et al.,* 2020). Mating pairs that produced larvae had the greatest impact on soil nutrients, likely through larval feeding and excretion distributing ammonium and nitrates into the soil (Parmenter and MacMahon, 2006; Hoback *et al*., 2020). However, previous studies considered only beetle presence, overlooking insect behaviour as a potentially important but poorly understood mechanism of carrion decomposition.

*Nicrophorus* beetles exhibit a remarkable range of complex parental care behaviours involved in preparing and maintaining carcasses for their broods (Figure 1). For instance, removing fur or feathers to increase access to labile tissues for offspring consumption (Potticary *et al*., 2024); burying their carrion brood balls belowground, increasing soil aeration (Brown *et al*., 2010); smearing antimicrobial exudates on the carcass surface (Duarte *et al*., 2018); and consuming and processing carrion tissue. The parental behaviour of beetles during this process therefore plays a crucial role in nutrient cycling, but it is also highly sensitive to environmental stressors. For instance, in two laboratory experiments using *N. nepalensis*, carcass preparation behaviour was negatively affected by warming. Specifically, the quality of carcass preparation decreased at elevated temperatures (Malik *et al*., 2025), despite a tendency for parents to spend more time preparing the carcass under warming (Malik *et al*., 2024). In *N. vespilloides,* a heatwave just before breeding increased male parental care duration (Pilakouta *et al*., 2023), and a developmental heatwave increased the likelihood of females providing care (Sidhu *et al*., 2024). These studies demonstrate the sensitivity of burying beetle behaviours to thermal stress, which suggests they may be particularly vulnerable to climate extremes. Understanding how climate-driven changes in their behaviour alter carrion decomposition is therefore essential for predicting nutrient cycling under future climate change.

**Figure 1.**
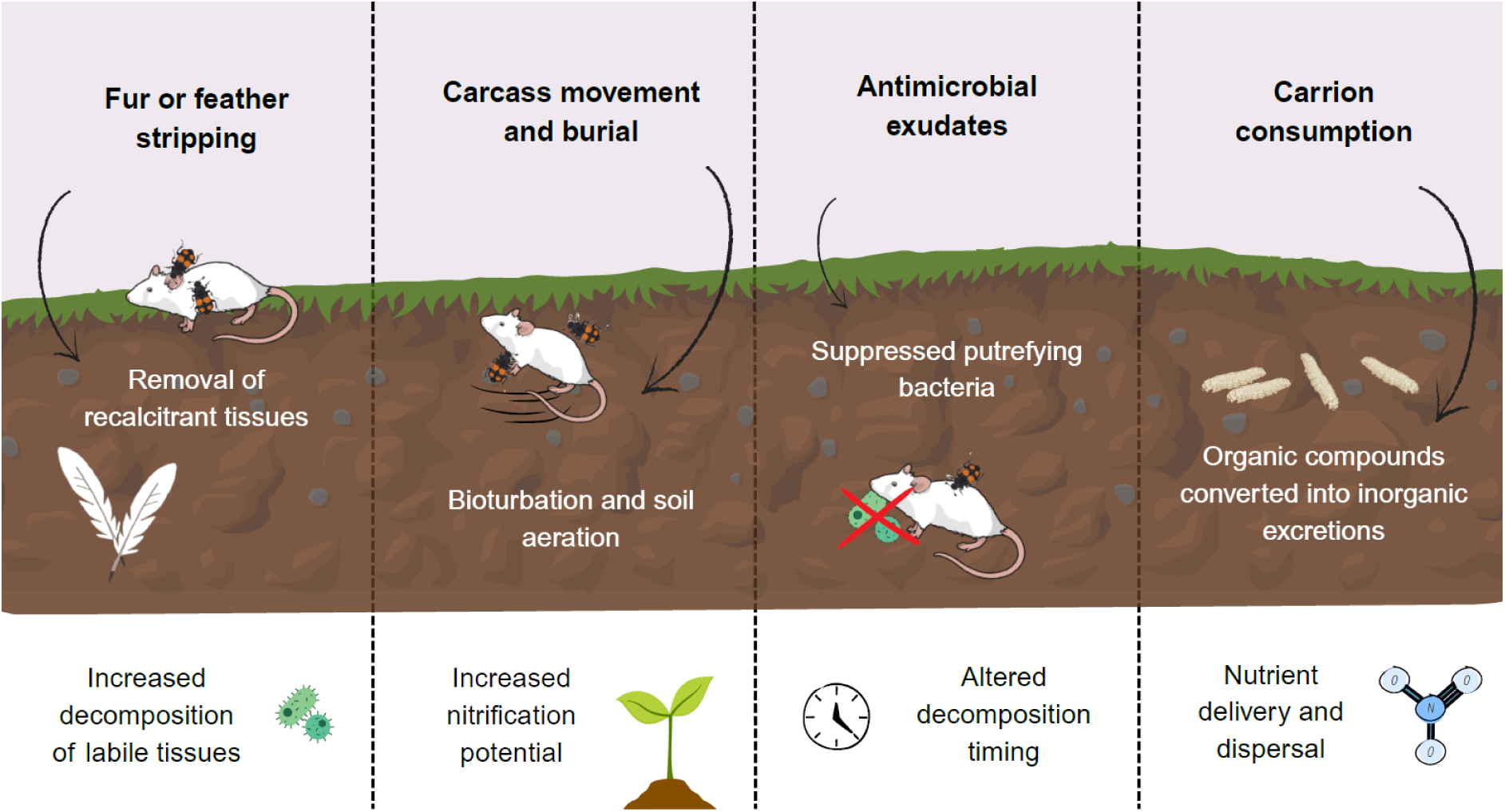
Conceptual illustration of burying beetle carcass preparation and hypothesised links to carcass decomposition and carrion-derived nutrient delivery. Fur and feathers contain keratin, a recalcitrant and insoluble protein that can constrain decomposition (Palacz-Ziółek *et al*., 2025). Carcass movement and burial could reduce soil compaction and promote nitrification (Manning *et al*., 2016). Chemical modification of the carcass surface could alter putrefying microbial communities (Duarte *et al*., 2018; Shukla *et al*., 2018), and carcass consumption and frass and urea deposition may facilitate the transfer of nutrients from carcass biomass into more bioavailable forms (Bartel *et al*., 2024). Beetle icons produced by Jakob Wiil.

In this study, we used *N. vespilloides* beetles to test whether heatwaves disrupt carrion decomposition and nutrient cycling due to changes in burying beetle behaviour. To address this question, we used a simulated heatwave and measured carcass decomposition and the input of carrion-derived nutrients to soil, both in the presence and absence of *N. vespilloides*. Crucially, we also examined potential behavioural and reproductive mechanisms that may underlie any changes in nutrient cycling, by measuring carcass preparation behaviour, parental care, and reproductive success. We predicted that beetle presence would enhance carcass decomposition, increasing the input of carrion-derived nutrients into the soil. We also hypothesised that heat stress would negatively impact this process, driven by heatwave-induced reductions in parental care resulting in impaired carcass preparation behaviour and reduced reproductive success. As climate extremes become more frequent, understanding how insect behavioural responses shape decomposition processes will be critical for predicting ecosystem functioning.

## Materials and Methods

### Study species

*Nicrophorus vespilloides* is a carrion beetle species with parental care, characterised by obligate carcass preparation and facultative parent-offspring interaction. After finding a carcass suitable for breeding, parents bury it in soil, remove any fur or feathers, and secrete antimicrobial exudates to reduce microbial competition. The female lays eggs in the surrounding soil, which hatch and crawl to the carcass approximately 2-3 days after oviposition. Larvae can be provisioned with pre-digested food by their parents but also self-feed before dispersing from the carcass to pupate in the soil, typically less than 1 week later. They eclose into adults around 21 days after dispersal and become sexually mature 10-14 days later (Scott, 1668; Potticary *et al*., 2024).

### Animal husbandry

We used fifth-generation lab-reared beetles maintained at the University of St Andrews in the UK. The laboratory population originated from wild beetles caught in Morton Lochs National Nature Reserve, Fife, Scotland in July and August 2025. Beetles were stored in individual transparent plastic containers (12 x 8 x 2 cm) filled with moist compost (Clover Grow, UK). They were kept at 20°C in cooled incubators (LMS Series IA Model 280 NP) in complete darkness to mimic subterranean conditions and fed cubes of organic beef *ad libitum* twice a week.

### Soil collection and microcosm assemblage

Soil was collected under license from Morton Lochs NNR due to the known population of *N. vespilloides* in this area and its previous successful use as a breeding substrate (Grieve and Pilakouta, 2026b). Five collection points were chosen at random within a native mixed broadleaf stand (1000m²) (Supplementary Figure 1). A steel corer (5cm diameter x 10cm depth) was used to extract topsoil, which was then homogenised through a 2mm sieve and stored at 4°C. Microcosms of transparent plastic containers (17 × 12 × 6 cm) were each filled with 150g of soil. A freshly thawed mouse carcass of a standard size (16-20g) (Livefoods Direct) was weighed using an Ohaus PR124 analytical balance (±0.0001 g) and added to each microcosm.

### Experimental design

Our experiment followed a 2 x 2 full factorial randomised block design, with two factors completely crossed and randomly assigned to 160 microcosms that contained soil and a mouse carcass. The two factors were thermal treatment (control or heatwave) and beetle presence (with or without beetles), resulting in four treatment combinations (n = 40 per treatment) that were stratified across four blocks.

The four treatments were (i) control treatment without beetles where microcosms were kept at 20°C throughout the experiment, (ii) control treatment with beetles where microcosms contained a mating pair and were kept at 20°C throughout the experiment, (iii) heatwave treatment without beetles, where microcosms were exposed to a 72hr simulated heatwave (26°C), and (iv) heatwave treatment with beetles, where microcosms contained a mating pair and were exposed to a 72h simulated heatwave (26°C). (Figure 2). The temperature and duration of the simulated heatwave has previously been shown to be stressful for reproduction and parental care in this species (Pilakouta *et al*., 2023; Grieve and Pilakouta, 2026a) and is comparable with predicted heatwaves in central Scotland (O’Neill and Tett, 2016). To avoid any effects of inbreeding, mating pairs were formed of virgin males and females that did not share any common ancestors for the past two generations. Beetles were mated within 20 days of sexual maturity (i.e., 10-30 days post-eclosion).

**Figure 2.**
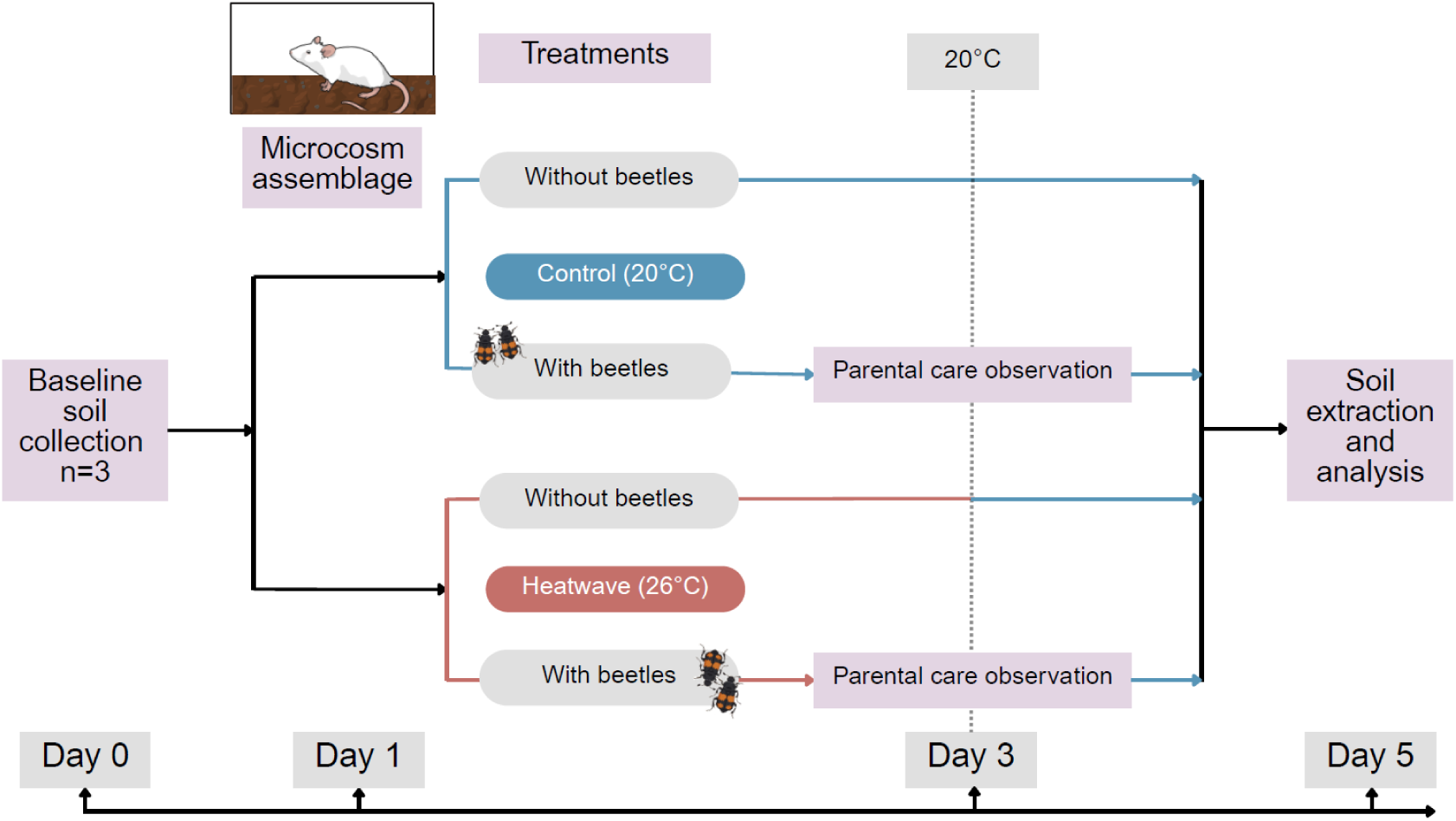
Experimental design with microcosms assigned one of two temperature treatments (control, 20°C; or heatwave, 26°C), with or without a mating pair of burying beetles. In microcosms with beetles, parental care behaviour and reproductive success was measured. Following the heatwave (72hrs), all microcosms were maintained at 20°C until the end of the experiment. Soil was collected from all microcosms on Day 5 for extraction and analyses.

Data for the four treatment groups were collected concurrently using two incubators set to constant temperatures (20°C and 26°C). The heatwave treatment microcosms were moved into the 20°C incubator after the simulated heatwave. Microcosms were maintained under experimental conditions for five days, after which the experiment was terminated and measurements were collected. This duration was selected to capture early-stage carcass decomposition and beetle behaviour within a standardised timeframe, whilst also aligning with the sampling timepoint used by Ilardi *et al*. (2021). Due to logistical constraints, the 160 microcosms were established in four sequential blocks over a four-week period (n = 40 per week). Treatments were balanced across blocks to ensure equal representation each week. Microcosms with no eggs (n = 5) were excluded from analyses as the likelihood of parental carcass preparation may have been influenced by the absence of developing offspring.

### Parental care assays

As we were interested in carcass preparation behaviour, and antimicrobial exudates peak on the third day after mating (Cotter *et al*., 2013), we conducted behavioural assays at 72h, prior to hatching (Figure 2). Before beginning the behavioural assays, the mating boxes were acclimated for 15 minutes under a red-light in a temperature-controlled room (20°C). We identified the male and female and recorded their behaviour using instantaneous scan sampling every minute for 30 minutes (developed by Dawkins *et al.,* 1664). Observed behaviours were grouped into three categories: carcass preparation (moving the carcass, antimicrobial smearing, fur stripping, crypt-making); other parental care behaviour (guarding the carcass), and non-parental behaviour (absence from the carcass, walking, self-grooming, self-feeding, or social interactions with partner).

To account for carcass preparation that may have occurred outside the observation window, we also gave the carcasses an ordinal score reflecting the preparation status at the end of the experiment (Supplementary Figure 2). This was based on three criteria: fur removal, formation of a crypt, and rolled into a ball. Carcasses received a score of 1 if all three criteria were fulfilled, 0.5 if one or two criteria were fulfilled, and 0 if there was no preparation (adapted from Duarte *et al*., 2018; Malik *et al*., 2025).

### Reproductive success and carrion consumption

The number of eggs laid was recorded by checking the underside of each beetle microcosm 24h after mating was initiated and twice daily thereafter. This method is minimally disruptive, and there is a strong positive correlation with the total number of eggs laid (Monteith, Andrews and Smiseth, 2012; Bladon *et al*., 2020). The number of unhatched eggs was counted at the end of the experiment to give hatching success as the proportion of total eggs that hatched. The number of larvae (brood size) was counted five days after carcass deposition, which also provided data on post-hatching survival when calculated as a proportion of hatched eggs. We also recorded the total mass of the larvae (brood mass) to estimate mean offspring size by dividing it by the number of larvae. The male and female parents were weighed before and after the experimental period to quantify mass change during breeding, calculated as the proportional difference. Caregiving adults gain mass through feeding on the brood ball (Gray *et al*., 2018), so mass change was used as a measure of carrion consumption (Pilakouta *et al*., 2016).

### Carcass decomposition

Carcass decomposition was quantified as the change in carcass mass over the five-day experimental period, a commonly used proxy for decomposition rates (e.g., Barton and Evans, 2017; Iyer *et al*., 2025). Each carcass was weighed prior to deposition in the microcosm. After the five days, the carcasses were removed, gently cleared of adhering soil, and reweighed to obtain final mass and the proportional mass loss. The exponential decay rate was calculated to express decomposition as a mass and time-standardised number, enabling a descriptive comparison across all types of organic matter. The exponential decay rate constant (k) was calculated for each carcass assuming a negative exponential decay model:

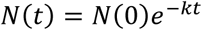

where *N*_0_is the initial carcass mass, *N_t_* is the mass at time *t* (5 days). Rearranging terms gives us the following equation: 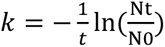

### Soil sampling and extractions

After carcass and beetle removal, the soil from each microcosm was individually collected and homogenised again through a 2mm sieve. To verify that soil properties remained stable over time, three baseline soil samples were collected each week and analysed alongside the experimental samples (Figure 2). Dissolved organic carbon (DOC) and dissolved organic nitrogen (DON) were measured by extracting 5g (fresh weight) of soil in 35mL ultrapure (Milli-Q*^®^*) H_2_O and analysed using an Elementar EnviroTOC analyser (Langenselbold, Germany). A further 5g of soil was extracted with the same method to measure dissolved inorganic N (NH₄⁺, and NO_3_^−^) using a Seal Analytical AQ400 discrete analyser (Mequon, WI, USA). All extracts were filtered through ashless Whatman No. 42 filter papers and syringe filtered to 0.45μm pore size. Soil pH (1:2 soil:CaCl_2_) was determined using a pH meter (Mettler Toledo, UK) and soil moisture content (SMC) was determined gravimetrically after drying at 105°C for 48hrs. Sample sizes for DOC, NH₄⁺, NO_3_^−^, pH and SMC measurements were as follows: n=38 for control treatments without a mating pair of beetles, n=38 for heat-exposed treatments without a mating pair of beetles, n=36 for control treatments with a mating pair of beetles, and n=38 for heat-exposed treatments with a mating pair of beetles. The second and third blocks of the dissolved organic nitrogen samples were removed from subsequent analyses due to instrument failure when measuring total N. Final sample sizes for DON therefore ranged from n=15-21 per treatment.

### Statistical analysis

We conducted our analyses in three stages to progressively refine inference from treatment-level effects to underlying mechanisms. Initial factorial models tested our first hypothesis that beetle presence enhances carcass decomposition and ecosystem responses, and our second hypothesis that these responses are reduced under heat stress. Subsequent models tested whether heat stress affected the beetle traits hypothesised to underpin these effects (carcass preparation behaviour and reproductive success). Finally, piecewise structural equation models integrated these relationships to assess direct and indirect causal pathways. All statistical analyses were conducted using R (version 4.5.2) (R Core Team, 2025) and figures were generated using ‘ggplot2’ (version 3.5.2) (Wickham *et al*., 2026). Significance of the fixed effects was assessed using Type II Wald chi-square tests using the ‘Anova’ function from the ‘car’ package (Fox and Wesiberg, 2016).

#### (i) Factorial models

The proportion of carcass mass lost was analysed using a beta-distributed generalised linear mixed-effects model from the ‘glmmTMB’ package (Brooks *et al.,* 2026). Linear mixed-effects models were fitted using the ‘lme4’ package (Bates *et al*., 2015) for the soil response variables. Temperature stress, beetle presence, and their interaction were included as fixed effects, with block fitted as a random intercept. Carcass mass can influence soil nutrient input (Jones *et al*., 2025) and was considered as a covariate in all soil models. Model assumptions were evaluated using simulation-based residual diagnostics implemented using the ‘DHARMa’ package (Hartig, 2024). Covariate selection was based on AIC, as described in the Supplementary Methods, with final model specifications provided in Supplementary Table 1.

#### (ii) Effect of temperature stress on beetle behaviour and reproduction

Carcass preparation behaviour was analysed using a hurdle model from the ‘pscl’ package (Zeileis *et al*., 2008) as there was a high proportion of zeros in the data. The model simultaneously tested the likelihood of an occurrence and the amount of the behaviour, using binomial and negative-binomial components, respectively. Carcass preparation score was analysed using a cumulative link model (ordinal logistic regression) fitted with the clm function from the ‘ordinal’ package (Christensen, 2025). Temperature treatment was the main explanatory variable, with block and carcass mass as covariates. Male and female prebreeding mass were also considered as parental mass can influence temperature tolerance in this species (Grieve and Pilakouta, 2026c). Covariate selection was based on AIC and is described in Supplementary Methods.

Male and female mass change were analysed using Gaussian linear mixed-effects models, with temperature treatment as the primary explanatory variable and block as a random effect. Reproductive traits were analysed using generalised linear mixed-effects models with appropriate error distributions (Supplementary Table 1).

#### (iii) Path analysis

To examine the direct effects of heat stress on decomposition and soil nutrient inputs, and indirect effects mediated by burying beetle behaviour and reproductive success, we used piecewise structural equation modelling (pSEM) to test hypothesised causal pathways among variables. Models were implemented using the ‘piecewiseSEM’ package (Lefcheck, 2016), which provides flexibility for non-normal error distributions and mixed model types.

Analyses were restricted to beetle-only treatments. To maintain a consistent sample size across models, we only included observations with complete data for all variables (n=72). Heat stress was our exogenous variable and given the number of potential behavioural and reproductive predictors, we reduced model complexity *a priori*. Specifically, we retained a single measure of carcass preparation behaviour (total behaviour duration) as both measures were significantly correlated (*r* = 0.34, *p* = 0.002). Brood size was included as the primary reproductive trait as it is a key determinant of carcass decomposition (Hoback *et al.,* 2020). In burying beetles, parents actively regulate brood size through infanticide when offspring numbers exceed what the carcass size can support (Bartlett, 1687; Schrader *et al*., 2015; Hsu *et al*., 2024). Male and female mass change were included as measures of parental consumption, and carcass mass loss was incorporated as an intermediate pathway linking beetle activity to soil nutrient inputs. The same general model structure was applied to all soil response variables (DOC, NH₄⁺, NO_3_^−^, pH and soil moisture content), with the exception of dissolved organic nitrogen, which was excluded due to an insufficient sample size. This approach, using five separate but structurally consistent models, avoided over-parameterisation of the residual correlation structure among soil variables relative to the available data.

Model fit was assessed using tests of directed separation and Fisher’s C statistic, with the conservative option used when evaluating conditional independence claims (Supplementary Table 2). Block was retained as a random effect in component models where variance estimates supported its inclusion and omitted when variance was effectively zero to avoid singular or over-parameterised models. Zero-adjusted models (total maintenance behaviour and brood size) are not currently supported within the pSEM framework; therefore, negative binomial models were used for these responses to maintain compatibility, acknowledging that this approach does not explicitly account for zero inflation. Standardised path coefficients were calculated to enable comparison of effect sizes across pathways, but coefficients from negative binomial models were interpreted on the log-link scale.

## Results

### Carcass decomposition

Beetles increased carcass mass loss, but this effect was reduced under heat stress (heat stress: LRχ² = 5.65, p = 0.015; beetle presence: LRχ² = 52.27, p < 0.001; interaction: LRχ² = 11.90, p < 0.001, Figure 3). Mean carcass decay rate constants (k) were 0.0037 day⁻¹ for control treatments without beetles, 0.0176 day⁻¹ for control treatments with beetles, 0.0054 day⁻¹ for heat treatments without beetles, and 0.0063 day⁻¹ for heat treatments with beetles.

**Figure 3.**
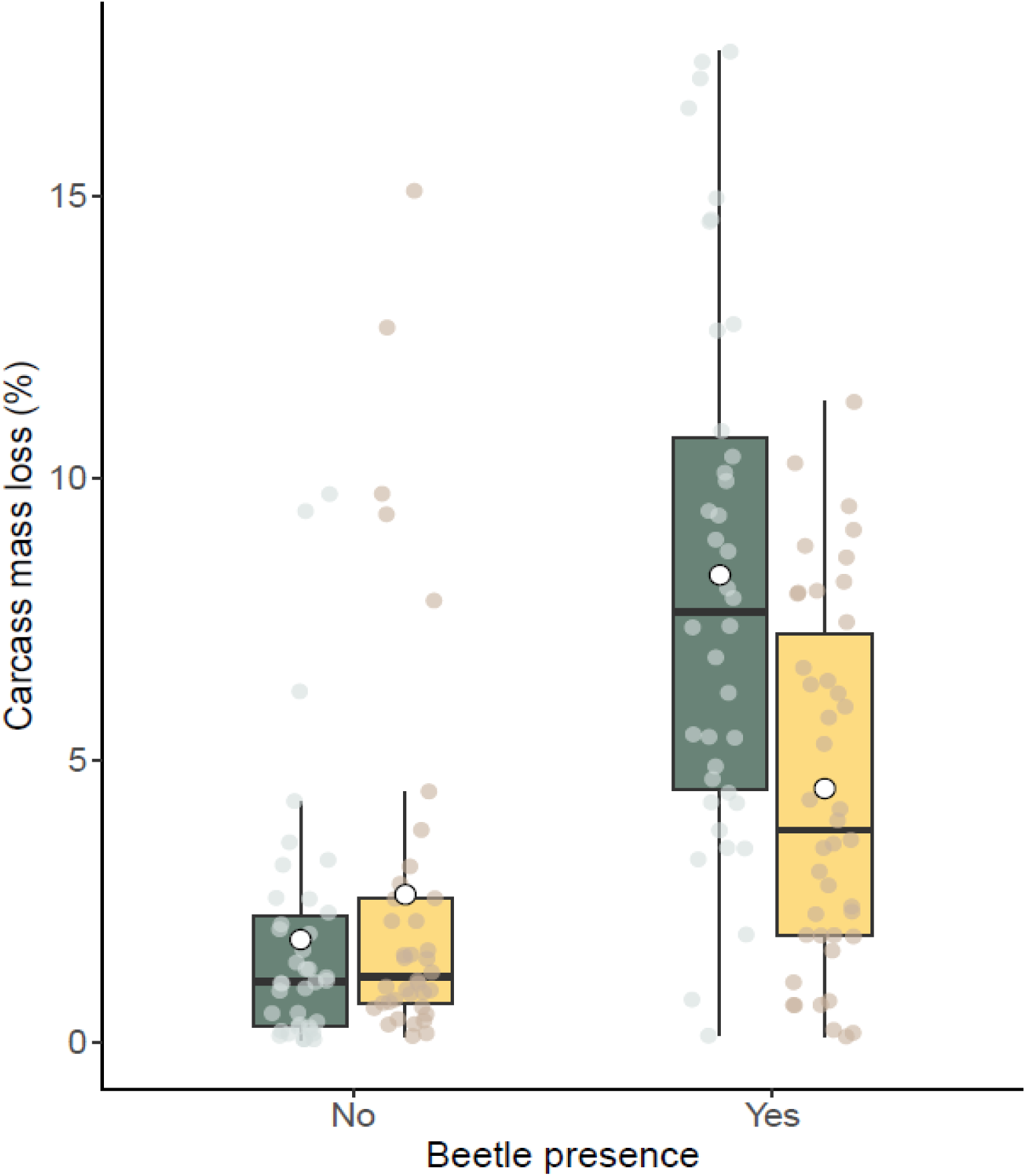
Interactive effects of beetle presence and heat stress on carcass mass loss. Carcass mass at day five as a percentage of starting mass for carcasses with and without a mating pair of burying beetles under control (green) and heatwave (yellow) treatments. Boxplots display a median line, interquartile range boxes, 1.5 x IQR whiskers and individual data points. White circles represent the mean values.

### Soil nutrients

Soil dissolved organic carbon (DOC), ammonium (NH_4_^+^), pH and moisture all significantly increased under heat stress and in the presence of beetles (Table 1; Figure 4A,C,E,F), whilst nitrate (NO_3_^-^) only significantly increased in beetle presence (Table 1; Figure 4D). In contrast, dissolved organic nitrogen (DON) did not change across treatments (Table 1, Figure 4B) and for no measure of soil nutrients was there an interaction between heat stress and beetle presence, though this was marginally nonsignificant for soil moisture (Table 1).

**Figure 4.**
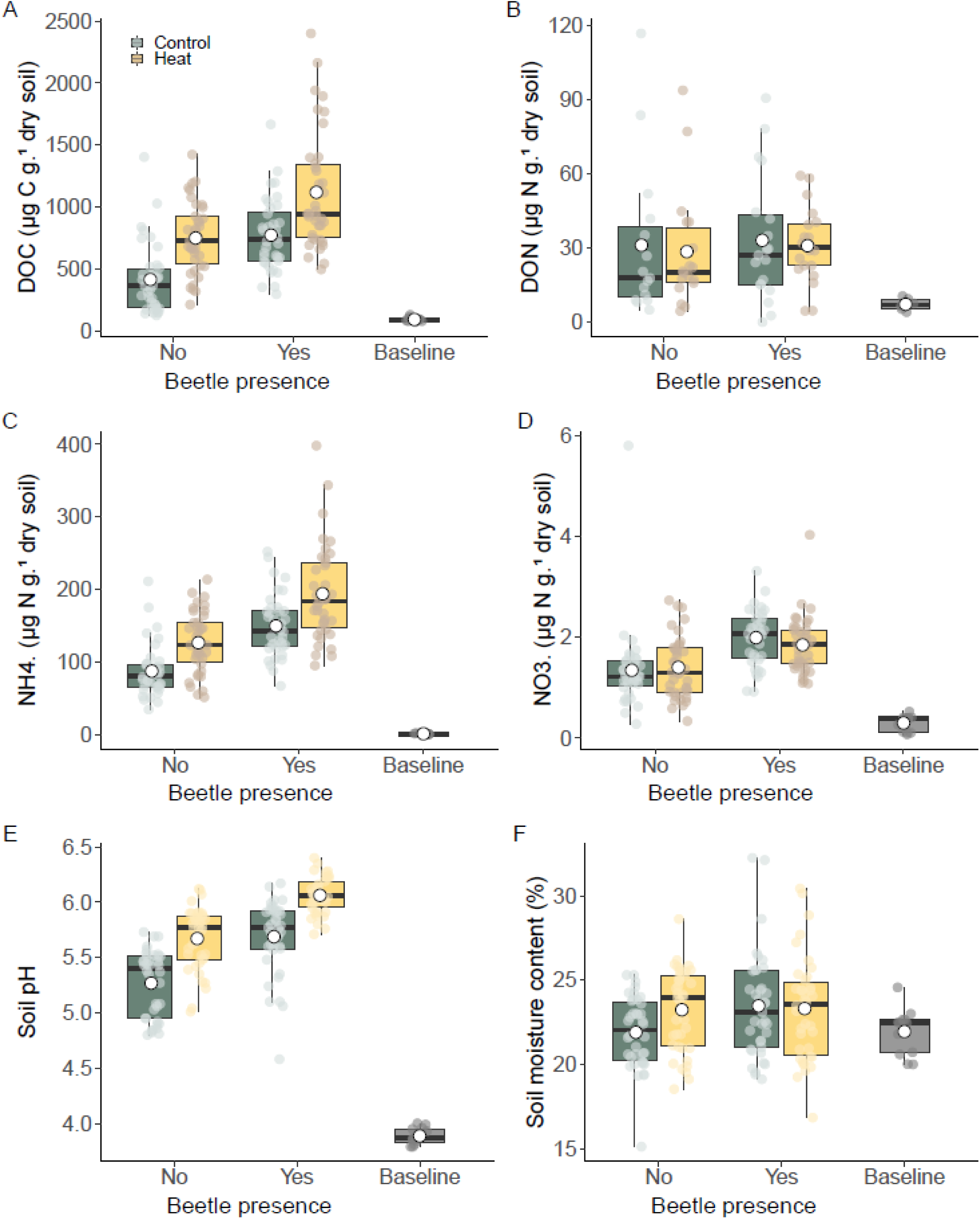
Effects of beetle presence and heat stress on soil properties. Boxplots show (A) dissolved organic carbon (DOC), (B) dissolved organic nitrogen (DON), (C) ammonium (NH₄⁺), (D) nitrate (NO₃⁻), (E) soil pH, and (F) soil moisture content under control and heat-stress treatments in the presence or absence of beetles. Boxplots display a median line, interquartile range boxes, 1.5 x IQR whiskers, and individual data points. The baseline consisted of three independent replicates of soil freshly collected and processed before each block (n=12).

**Table 1.**
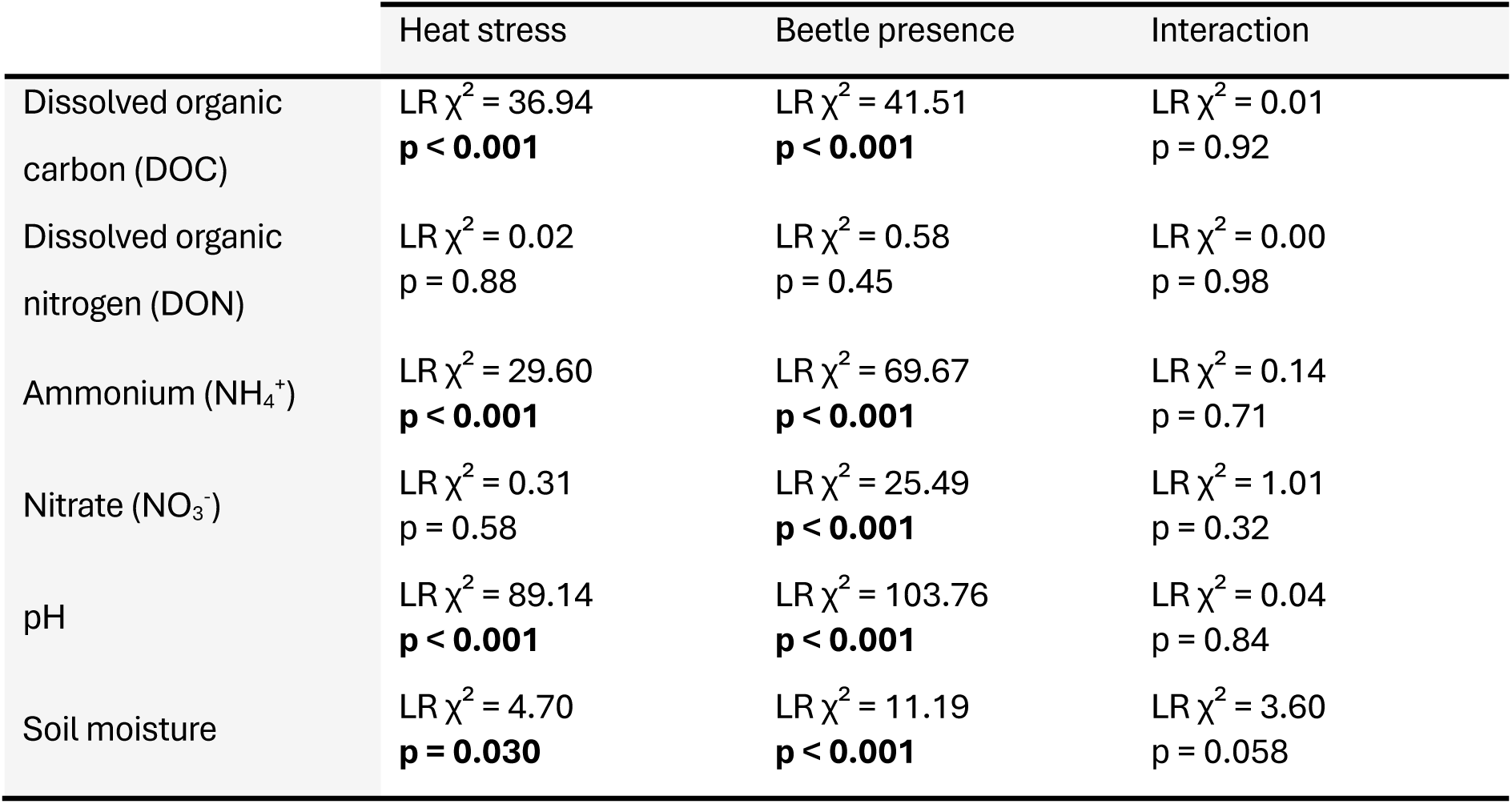
Results of likelihood ratio tests for main and interactive effects of heat stress and beetle presence on soil nutrients, pH and moisture. Statistically significant p-values are indicated in bold.

|  | Heat stress | Beetle presence | Interaction |
| --- | --- | --- | --- |
| Dissolved organic carbon (DOC) | LR $\chi^2 = 36.94$<br>**$p < 0.001$** | LR $\chi^2 = 41.51$<br>**$p < 0.001$** | LR $\chi^2 = 0.01$<br>$p = 0.92$ |
| Dissolved organic nitrogen (DON) | LR $\chi^2 = 0.02$<br>$p = 0.88$ | LR $\chi^2 = 0.58$<br>$p = 0.45$ | LR $\chi^2 = 0.00$<br>$p = 0.98$ |
| Ammonium ($\text{NH}_4^+$) | LR $\chi^2 = 29.60$<br>**$p < 0.001$** | LR $\chi^2 = 69.67$<br>**$p < 0.001$** | LR $\chi^2 = 0.14$<br>$p = 0.71$ |
| Nitrate ($\text{NO}_3^-$) | LR $\chi^2 = 0.31$<br>$p = 0.58$ | LR $\chi^2 = 25.49$<br>**$p < 0.001$** | LR $\chi^2 = 1.01$<br>$p = 0.32$ |
| pH | LR $\chi^2 = 89.14$<br>**$p < 0.001$** | LR $\chi^2 = 103.76$<br>**$p < 0.001$** | LR $\chi^2 = 0.04$<br>$p = 0.84$ |
| Soil moisture | LR $\chi^2 = 4.70$<br>**$p = 0.030$** | LR $\chi^2 = 11.19$<br>**$p < 0.001$** | LR $\chi^2 = 3.60$<br>$p = 0.058$ |

### Parental care

The simulated heatwave decreased both the likelihood of carcass preparation behaviour (z = -3.60, p < 0.001, Figure 5A) and the duration of this behaviour during the observation window (z = -3.46, p < 0.001, Figure 5B). Inspection of Pearson residuals identified one observation with a relatively large residual, but sensitivity analyses showed that its exclusion did not affect the results (z = -3.46, p < 0.001), so all observations were retained.

**Figure 5.**
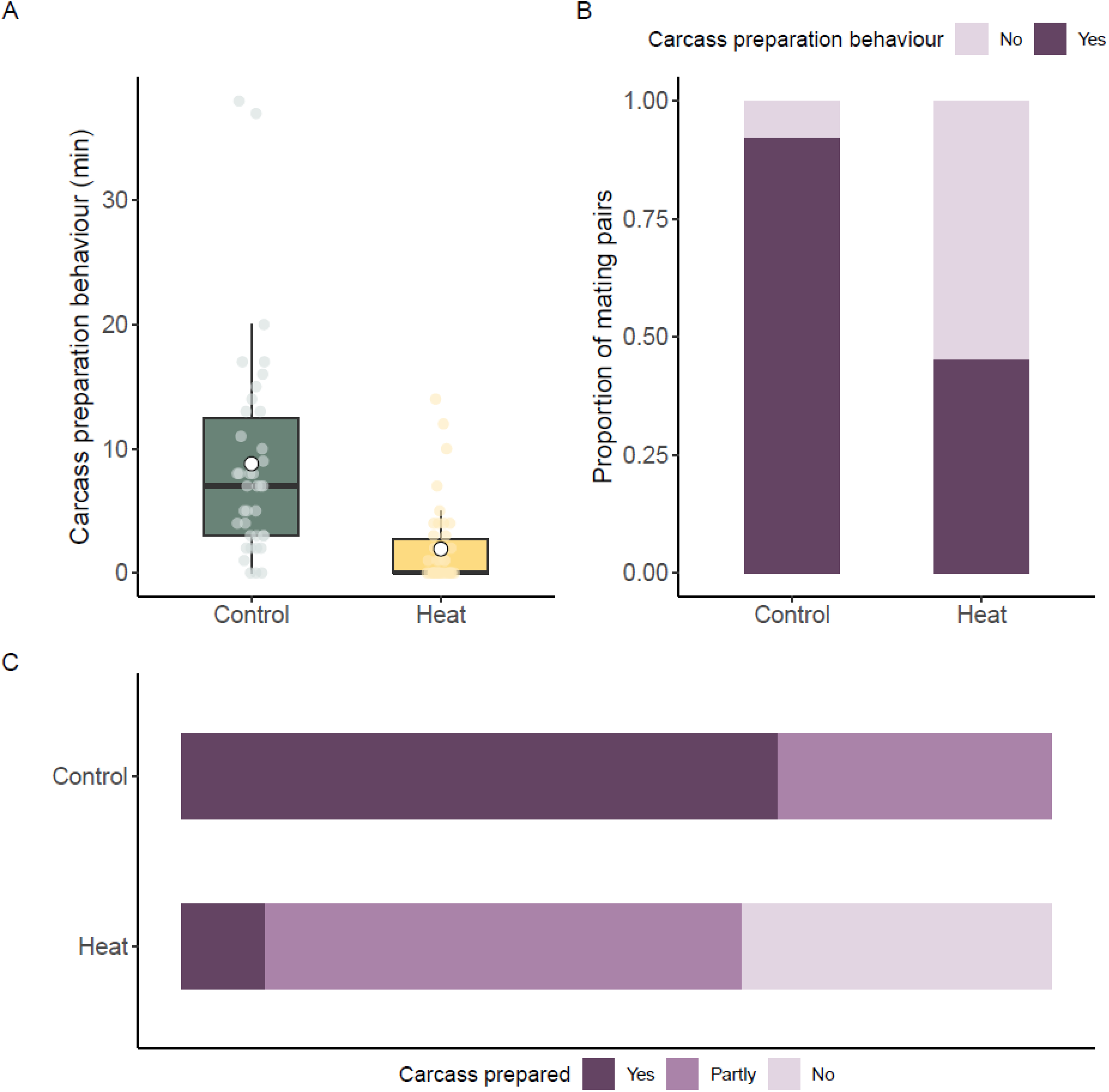
**Effects of heat stress on beetle carcass preparation behaviour**. (A) The amount of care (minutes) provided during the behaviour observation period for both male and female parent (60 minutes combined), (B) the proportion of mating pairs providing at least one instance of carcass preparation behaviour during the 30-min behavioural observation, and (C) the proportion of carcasses given an ordinal score of 0 (no preparation), 0.5 (partly prepared) and 1 (fully prepared).

Heat stress also influenced the total preparedness of the carcass as recorded by ordinal score (LRχ² = 27.37, p < 0.001). Fewer carcasses received a score of 1 (fully prepared) in the mating pairs under heat stress (Figure 5C). There was no difference in female or male mass change between control and heat treatments (LRχ² = 0.26, p = 0.56; LRχ² = 1.85, p = 0.17, respectively).

### Reproductive success and offspring fitness

We found that all measures of reproductive success were negatively affected by the heatwave. Fecundity (LRχ² = 13.25, p < 0.001), likelihood of producing offspring (z = -4.34, p < 0.001), number of offspring (brood size; z = -7.06, p < 0.001), and total brood mass (LRχ² = 56.16, p < 0.001) were significantly reduced in mating pairs exposed to the heatwave. All measures of offspring fitness were also reduced under heat stress (hatching success: LRχ² = 47.39, p < 0.001; post-hatching survival: LRχ² = 26.36, p < 0.001; average larval mass: LRχ² = 13.09, p < 0.001) (Figure 6).

**Figure 6.**
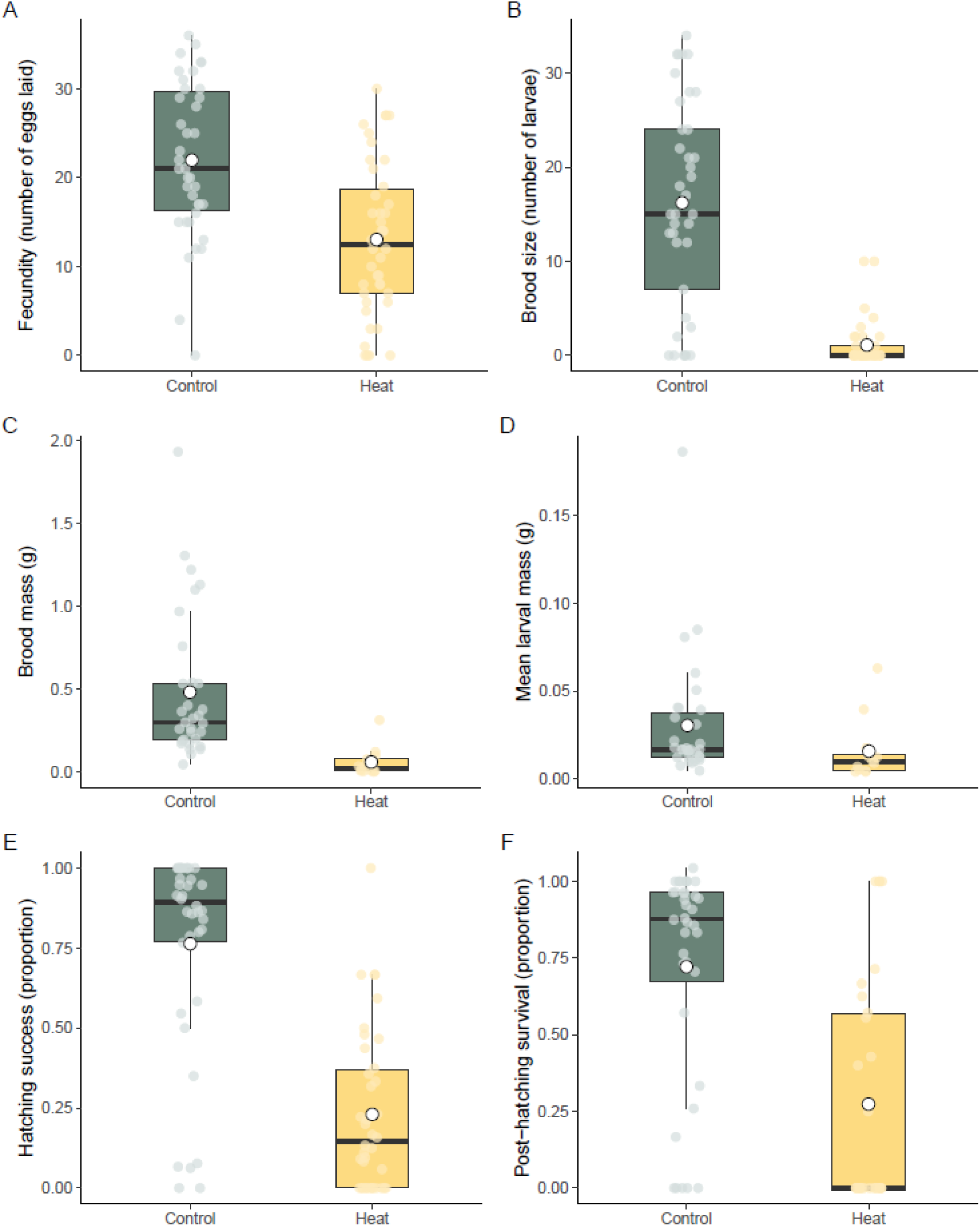
**Effects of heat stress on reproductive success and offspring fitness**. (A) Fecundity (number of eggs laid), (B) brood size (number of offspring on the carcass at day 5), (C) total brood mass at day 5 (g), (D) mean larval mass at day 5 (g), (E) hatching success, and (F) post-hatching survival. Boxplots display a median line, interquartile range boxes, 1.5 × IQR whiskers, and individual data points. White circles represent the mean values.

### Path analysis

The piecewise SEMs were consistent with the hypothesised causal structure (Fisher’s C = 6.895, p = 0.548). The structural equation models explained substantial variation in brood size (R²_mar_ = 0.67), carcass preparation behaviour (R²_mar_ = 0.36), carcass mass loss (R²_mar_ = 0.34), soil pH (R²_mar_ = 0.38) and soil moisture content (R²_mar_ = 0.30), whilst a smaller proportion of variation was explained for dissolved organic carbon (R²_mar_ = 0.27), ammonium (R²_mar_ = 0.24), and nitrates (R²_mar_ = 0.14). Temperature stress explained little to no variation in female mass change (R²_mar_ < 0.001) or male mass change (R²_mar_ = 0.006).

The pSEMs identified direct relationships between the temperature treatment and three variables, with heat stress increasing soil pH, but reducing brood size and carcass preparation behaviour (Figure 7). In turn, brood size increased carcass mass loss, and carcass mass loss was associated with increased soil moisture content. Carcass preparation behaviour increased both soil moisture and nitrate concentrations whereas male mass change was linked to higher dissolved organic carbon and ammonium. Lastly, increased male mass gain was associated with greater carcass mass loss, although this was marginally non-significant (p = 0.061).

**Figure 7.**
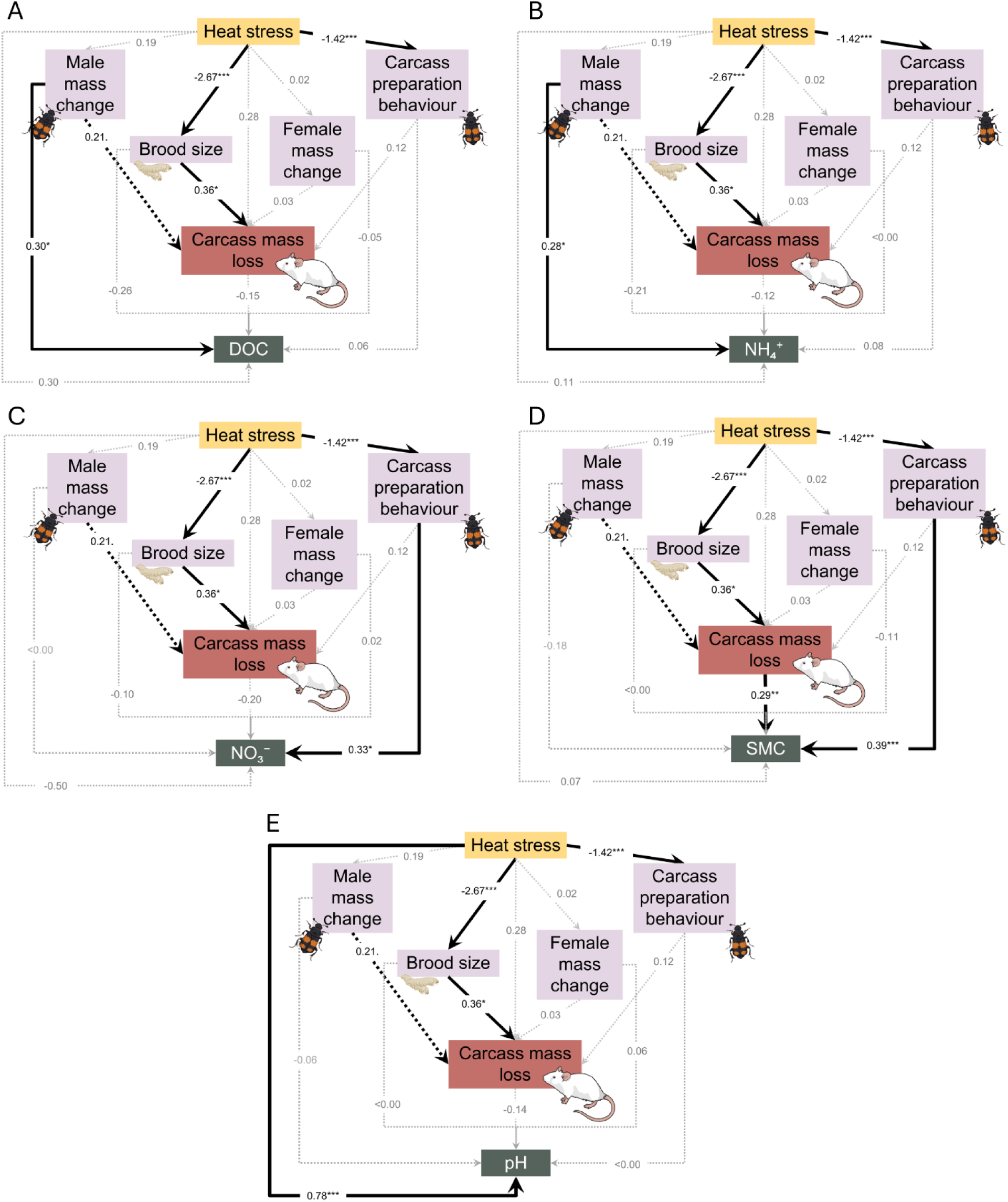
**Path diagrams explaining heat stress and beetle effects on soil**. (A) Dissolved organic carbon, (B) ammonium, (C) nitrates, (D) soil moisture content, and (E) pH. Path coefficients are given on top of the arrows. Coefficients are standardized for Gaussian models, coefficients for negative binomial models (brood size and carcass preparation behaviour) are shown on the log-link scale, with positive coefficients representing an increase in the response variable, and negative coefficients showing a decrease. Nonsignificant paths are grey, marginally nonsignificant paths black and dashed, and significant paths are black and solid.

## Discussion

Climate extremes associated with global warming are altering ecosystem processes, yet their effects on carrion-driven nutrient cycling remain unknown. Our results show for the first time that heatwaves, which are increasing in frequency and severity due to climate change, can have strong indirect effects on carrion-derived soil nutrient availability moderated by changes in insect behaviour. Moreover, we show that these indirect effects on soil nutrient availability can be stronger than direct effects of higher temperatures, suggesting that shifts in insect behaviour are a crucial regulator of nutrient cycling under future climate extremes.

Our simulated heatwave substantially altered carrion decomposition dynamics, resulting in greater carcass mass loss when beetles were absent, as well as increased concentrations of dissolved organic carbon, ammonium, soil moisture and pH. Increased temperatures are known to enhance decomposition rates in carrion, with elevated enzymatic activity accelerating soft-tissue degradation (Sessa *et al*., 2026), such as in sheep carcasses (*Ovis aries*; Carter and Tibbett, 2006), rats (*Rattus rattus*; Carter *et al*., 2008), deer (*Cervus nippon;* Hachiya *et al*., 2026), and freshwater fish (Boros *et al*., 2025). Decomposing microbes proliferate under elevated temperatures (Conant *et al*., 2004; Carter and Tibbett, 2006; Pechal *et al*., 2013) and have higher metabolic activity (Dilly and Munch, 1666). This enhanced microbial decomposition can accelerate the release of carcass-derived fluids (Keenan *et al*., 2018; DeBruyn *et al*., 2025), potentially contributing to the greater carcass mass loss and elevated soil moisture observed under heat stress. These fluids typically contain carbon-rich fatty acids and polysaccharides, and nitrogen-containing proteins, amino acids, and biogenic amines such as cadaverine and putrescine (DeBruyn *et al*. 2025). Increased microbial degradation of these nitrogenic compounds can contribute to nitrogen mineralisation (Macdonald *et al.,* 2014), potentially explaining the increase in soil ammonium observed under temperature stress. Indeed, increased temperatures resulted in higher microbial turnover, which raised the accumulation of nitrogen in soil (Wang *et al*., 2020). The input of these alkaline putrefaction products, and the increase in microbial decomposition under heat stress, likely contributed to the higher pH values also detected in our heat treatments (Macdonald *et al*., 2014; Keenan *et al*., 2018; Rietz *et al*., 2025). Notably, the effects of heat stress were not consistent across all forms of nitrogen. Whilst ammonium concentrations increased significantly under heat stress, dissolved organic nitrogen and nitrates remained unchanged. The combination of increased soil moisture and augmented microbial activity depleting soil oxygen may have inhibited nitrifying bacteria (Fiedler *et al*., 2023; Das *et al*., 2025). Furthermore, the rate of dissolved organic nitrogen production via cell lysis and tissue breakdown may have balanced with an increased rate of microbial consumption and ammonification, keeping the net measurable DON pool relatively similar (Macdonald *et al*., 2014; Zainul Kamal *et al*., 2022).

Carcasses with *N. vespilloides* present also had increased mass loss, consistent with other studies investigating the effect of insect presence on carcass decomposition rates (e.g., Englmeier *et al*., 2023; Iyer *et al*., 2025). Beetle presence was associated with elevated DOC, NH₄⁺, and NO₃⁻ concentrations, as well as increased soil pH. These findings support those of Ilardi *et al.,* (2021), who reported increases in soil organic matter and pH in the presence of breeding *N. vespilloides.* They are also in line with the results of Hoback *et al.,* (2020), who found elevated pH and nitrate concentrations in soils associated with breeding *N. marginatus.* Interestingly, we also detected an interaction between beetle presence and heat stress for carcass mass loss and soil moisture content, with heat stress reducing the magnitude of beetle effects on both variables. This finding suggests that warming altered aspects of burying beetle behaviour or reproductive performance that contribute to carrion decomposition and associated soil responses.

To investigate these pathways, we examined the effects of heat stress on carcass preparation behaviour, reproduction and parental mass change. The simulated heatwave decreased parental care, as measured by carcass preparation behaviour observed during the observation window and by carcass preparation score at the end of the experiment. This reduction in carcass preparation behaviour under heat stress is in line with other experiments using *Nicrophorus* spp. (Malik et al. , 2024; Pilakouta *et al*., 2024; Malik *et al*., 2025) and may reflect the cost of self-maintenance and survival exceeding the net fitness benefit of parental care for offspring survival (Grew *et al*., 2016). For instance, heat stress is associated with higher metabolic costs and increased oxidative stress during reproductive investment, resulting in a reduced capacity to provide care (Merkling *et al*., 2017). Parental mass change, as a proxy for somatic investment was not affected by exposure to the heatwave.

All of the reproductive success traits (fecundity, brood size and brood mass) and offspring fitness traits (hatching success, post-hatching survival, larval mass) that we measured were reduced under heat stress. Reproduction has been shown to be highly sensitive to heat stress across a range of taxa (e.g., Canal Domenech and Fricke, 2022; Chatten *et al*., 2025). Indeed, a similar heatwave of 72h during breeding was found to be particularly detrimental for reproduction in the same *Nicrophorus* species (Pilakouta *et al*., 2023). Increased temperature may have influenced mating behaviour (Pilakouta and Baillet, 2022) or gamete quality or quantity (Sales, Vasudeva and Gage, 2021). Lower gamete quality can also have downstream consequences for offspring (Alavioon *et al*., 2016), but our findings of decreased offspring fitness under heat stress may also be related to the observed reduction in parental care (Eggert, Reinking and Müller, 1668; Grew *et al*., 2016) or accelerated carcass decomposition and reduced resource acquisition (Arce *et al*., 2012).

Our pSEMs also identified relationships between beetle traits and carcass decomposition, which highlight the consequences of burying beetle reproductive and behavioural changes under heat stress. Greater carcass mass loss was linked with larger broods and was weakly associated with increased male mass gain. This may indicate increased carcass consumption, with more biomass conversion from carcass tissue into offspring and adult somatic mass (Parmenter and MacMahon, 2006; Pechal *et al*., 2013; Rupali *et al*., 2024). Carcass mass loss was also positively linked with soil moisture. As soil variables were measured after five days of decomposition, corresponding to the active decay stage in mice (Al-Mekhlafi *et al*., 2020), this relationship likely reflects the release of decomposition fluids as soft tissues liquefy. During active decay, the surrounding soil becomes increasingly saturated (DeBruyn, Keenan and Taylor, 2025), elevating soil moisture (Barton *et al*., 2016).

Although heat stress and beetle presence increased both carcass mass loss and soil carbon and nitrogen concentrations, pSEMs revealed no direct relationship between carcass mass loss and dissolved organic carbon, ammonium, or nitrate. This suggests that treatment effects on decomposition and nutrient dynamics may have occurred through parallel pathways rather than through a simple increase in carcass mass loss. Unlike soil moisture, which is closely linked to the release of decomposition fluids (Barton *et al*., 2016), soil carbon and nitrogen pools are shaped by multiple biological and biogeochemical processes, including microbial immobilisation (Raza *et al*., 2023) and bioassimilation (Parmenter and MacMahon, 2006). Furthermore, the soil nutrient concentrations represented a temporal snapshot of decomposition, whereas carcass mass loss reflected cumulative resource processing across the experimental period. Previous work has shown that the effects of burying beetles on soil organic matter can change over the course of decomposition (Ilardi *et al*., 2021), suggesting that variation in carcass mass loss may not necessarily correspond to nutrient concentrations measured at a single time point. Investigating these temporal dynamics was beyond the scope of the present study but represents an important direction for future work.

Male mass gain, on the other hand, was positively associated with dissolved organic carbon and ammonium concentrations. This is broadly consistent with increased carrion consumption promoting nutrient transfer to surrounding soils through the deposition of frass and excretions (Chen and Forschler, 2016; Tomberlin *et al*., 2017) or mechanical fragmentation of larger organic matter (Rupali *et al*., 2024). As females spend more time consuming carcass tissue and regurgitating it to offspring (Smiseth and Moore, 2002; Ratz *et al*., 2021; Ratz, 2025), more of the carrion they ingest may be allocated to offspring mass than the release of carbon and nitrogen into the soil, potentially explaining why female mass gain was not similarly associated with these nutrients.

One of the most notable findings from our pSEMs was that carcass preparation behaviour was positively associated with increased soil moisture and nitrate concentrations. Fur stripping may have removed the insulating carcass layer typically slowing dessication (e.g., Englmeier *et al*., 2023), and crypt-making could have allowed internal fluids and liquefied tissues to seep into the soil, increasing moisture. However, carcass preparation behaviour was not associated with carcass mass loss, which would be expected to increase if these behaviours were the primary mechanisms driving soil moisture. Instead, the preparation behaviour may have altered soil structure and facilitated fluid infiltration. For example, dung beetle tunnelling activity significantly increases steady-state soil infiltration rate (Keller *et al*., 2022), and soil foraging activity has been linked with increased water infiltration (Eldridge *et al*., 2010). This mixing and reworking of soils, or bioturbation, may also explain the increase in nitrate concentrations. Soil disturbance by animals can increase soil aeration and soil oxygen content (Pang *et al*., 2023), supporting the metabolism of aerobic microbes and promoting nitrification (Keenan *et al*., 2018). Furthermore, organic matter manipulated by soil foraging animals was found to be in closer contact with soil bacteria, which increased microbial decomposition and nutrient mineralisation (Eldridge and Mensinga, 2007) .

More generally, our findings of increased nitrate concentrations when beetles were present highlight the important role of burying beetles in regulating nutrient cycling during decomposition. Burying beetles can sequester and monopolise a substantial proportion of small vertebrate carrion; for example, Trumbo (2016) found that burying beetles monopolised over 75% of small vertebrate carcasses in a forest habitat, highlighting their important contribution to carrion decomposition in temperate forest ecosystems. Environmental change has also been associated with reductions in burying beetle abundance and diversity (Hoermann *et al.,* 2018), with implications for ecosystem functioning. Our findings further suggest that consequences for ecosystem processes may arise not only through changes in population size or community composition, but also through changes in behaviour. For instance, a reduction in brood size under warming resulted in more carcass mass remaining after five days of decomposition. Rapid microbial proliferation of the remaining biomass could result in a positive feedback loop, with climate-driven declines in beetles increasing microbial volatile emissions, including CO₂, CH₄ and N_2_O (Iyer *et al.,* 2025). The reduction in carcass preparation behaviour under warming could alter the rate and spatial distribution of nitrogen mineralisation, potentially changing the availability and timing of nutrients released into soils for uptake by plants. Such changes could have cascading consequences for soil nutrient availability and ecosystem productivity. This positive association between nitrate concentrations and carcass preparation behaviour represents just one example of how climate warming could alter ecosystem processes via behavioural changes. Understanding behavioural linkages between global climate change and ecosystem functioning therefore remains a crucial yet understudied aspect of predicting how ecosystems will respond to climate change.

## Conclusion

In sum, our findings suggest that whilst warming may directly alter carrion decomposition, sublethal changes in animal behaviour and reproduction can scale up to ecosystem processes and nutrient cycling. This highlights an overlooked mechanism by which climate extremes may affect ecosystem functioning. In particular, reductions in carcass preparation and brood size under simulated heat stress were associated with changes in carcass decomposition and nutrient availability. Given that carrion provides rapid nutrient fluxes that sustain a diverse array of organisms, warming-induced changes could have cascading effects on ecosystem productivity, stability and resilience. Importantly, the responses of ecosystem engineers such as burying beetles may either amplify or redirect these effects. Incorporating animal behaviour and reproduction will therefore be critical for predicting how increasingly frequent climate extremes affect the capacity of ecosystems to maintain essential functions in an uncertain future climate.

## Acknowledgments

We are very grateful to Ian Washbourne for his expert advice and extensive time spent analysing soil samples. We would also like to thank NatureScot for providing the licenses required to conduct this work, as well as Rebekah Douglas, Sam Wraight, Solène Morelle and Jakob Wiil for their assistance with beetle husbandry.

