## Supplementary Methods for "From individuals to ecosystems: carrion beetle responses to heatwaves impact nutrient cycling"

**Electronic Supplementary Material**

*Supplementary Methods*

Model assumptions were assessed using simulation-based residual diagnostics implemented in the DHARMa package (Hartig, 2024). These diagnostics were used to evaluate residual dispersion, distributional assumptions and other departures from model assumptions, and informed the choice of error distribution. Final model distributions and links are reported in Supplementary Table 1.

For models in which additional covariates were considered, we used stepwise model selection based on Akaike’s Information Criterion (AIC), implemented using the step function in the MASS package. Covariates were retained where their inclusion improved model fit (>2 AIC). For the factorial soil models, carcass mass was considered as a covariate because variation in carcass mass can influence nutrient inputs to soil (Jones *et al.*, 2025). For behavioural and reproductive models, carcass mass and parental prebreeding mass were considered. For male and female mass change, carcass mass and brood size were considered. Final model structures are provided in Supplementary Table 1.

*Supplementary Tables*

***Table S1.*** *Statistical model specifications for carcass decomposition, soil responses, beetle behaviour and reproductive traits.*

| **Response** | **Model** | **Distribution/response structure** | **Fixed effects** | **Random effects** | **Covariates** |
| --- | --- | --- | --- | --- | --- |
| **DOC** | LMM | Gaussian | Temperature x beetle presence | Block | Carcass mass |
| **DON** | LMM | Gaussian | Temperature x beetle presence | Block | Carcass mass |
| **NH₄⁺,** | LMM | Gaussian | Temperature x beetle presence | Block | Carcass mass |
| **NO_3_^−^** | LMM | Gaussian | Temperature x beetle presence | Block | - |
| **pH** | LMM | Gaussian | Temperature x beetle presence | Block | - |
| **SMC** | LMM | Gaussian | Temperature x beetle presence | Block | - |
| **Carcass mass loss** | GLMM | Beta/logit | Temperature x beetle presence | Block | - |
| **Preparation behaviour** | Hurdle | Binomial/negative binomial | Temperature | Block | - |
| **Preparation score** | CLM | Ordinal logit | Temperature | - | - |
| **Male mass change** | LMM | Gaussian | Temperature | Block | - |
| **Female mass change** | LMM | Gaussian | Temperature | Block | - |
| **Fecundity** | GLMM | Negative binomial | Temperature | Block | - |
| **Brood size** | Hurdle | Binomial/negative binomial | Temperature | - | - |
| **Brood mass** | GLMM | Gamma/log | Temperature | Block | - |
| **Hatching success** | GLMM | Beta/logit; cbind (hatched, unhatched) | Temperature | Block | - |
| **Post-hatching survival** | GLMM | Binomial; cbind (dead, survived) | Temperature | Block | - |
| **Offspring mass** | GLMM | Gamma/log | Temperature | Block | - |

***Table S2.*** *Results of directed separation tests used to evaluate conditional independence and potential non-independence among variables in the piecewise structural equation model.*

| Independence claim test | Critical value | p-value |
| --- | --- | --- |
| Brood size ~ Total carcass maintenance | -0.83 | 0.41 |
| Female mass change ~ Total carcass maintenance | -1.29 | 0.20 |
| Male mass change ~ Total carcass maintenance | 0.30 | 0.59 |
| Female mass change ~ Brood size | 0.44 | 0.66 |
| Male mass change ~ Brood size | 0.20 | 0.66 |
| Male mass change ~ Female mass change | 3.85 | 0.054 |

*Supplementary Figures*

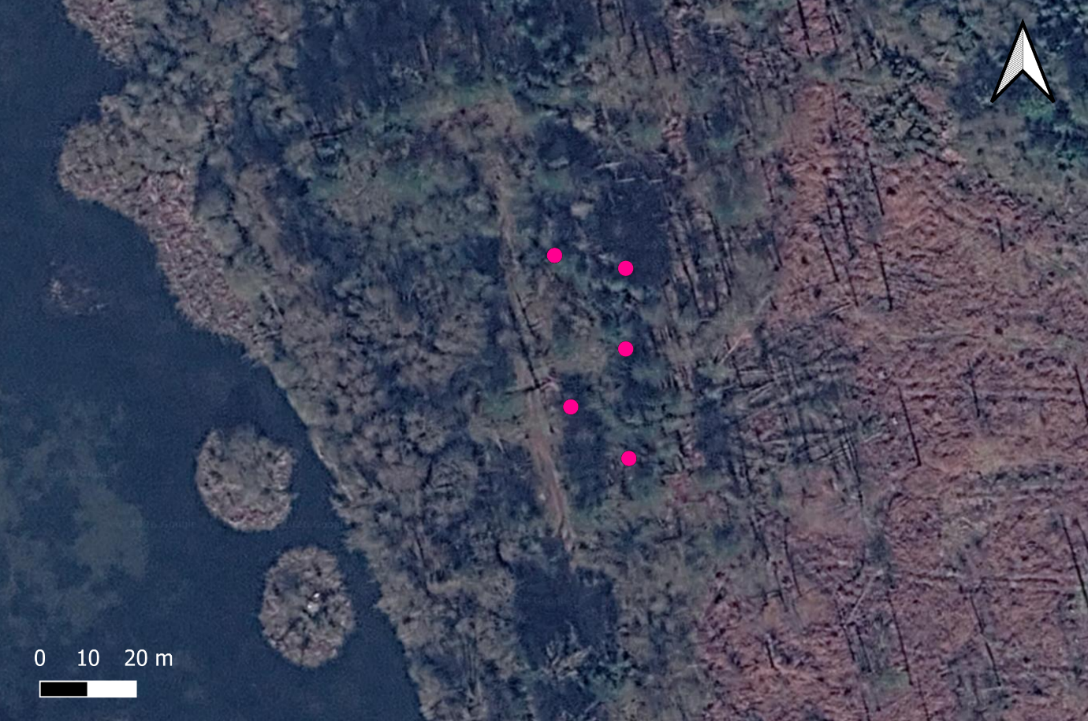

**Figure S1. Soil core sampling sites**. Cores were spaced 20m apart within a broadleaf stand in Morton Lochs Nature Reserve, Fife, Scotland.

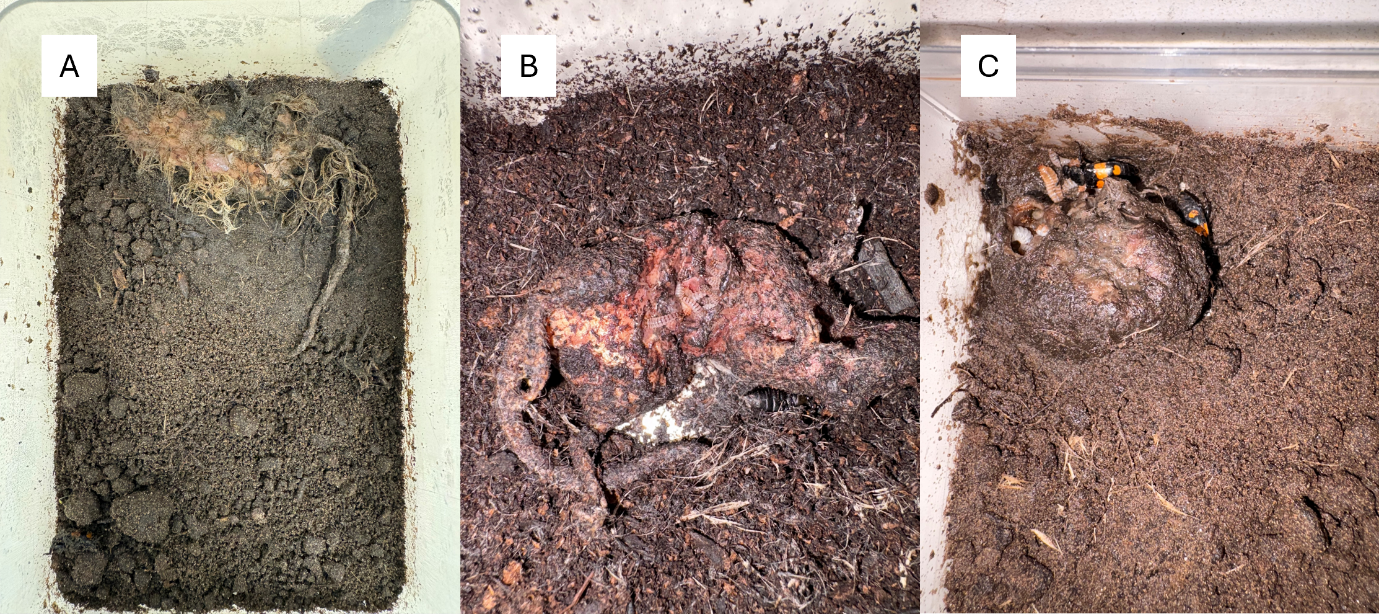

**Figure S2. Beetle-tended mouse carcasses on day five.** (A) Carcass given a preparation score of 0, as fur was not removed, there was no crypt, and the carcass had not been shaped into a ball. (B) Carcass scored as 0.5, as only one criterion was fulfilled (fur removal), and (C) scored 1 as there was fur removal, the carcass had a crypt, and was rolled into a ball.
